# Preventing injuries: How augmented reality reduces fall risks in pseudo-natural environments

**DOI:** 10.1101/2023.10.17.562672

**Authors:** F. Ball, J. Bomba, A. Nentwich, F. Meier, P. Vavra, T. Noesselt

**Author notes:** Correspondence to, Department of Biological Psychology, Otto-von-Guericke-University, PO Box 4120, 39106 Magdeburg, Germany. Shared first authors (F. Ball, J. Bomba).

## Abstract

Falling is a major contributor to injury-related incidents and the 2^nd^ most likely cause for unintentional injury deaths world-wide. Here, we therefore tested a vision aid for obstacle detection during a navigation task in pseudo-natural environments (virtually created cities). The vision aid had nine different setups (i.e. no highlighting of hazardous objects vs. eight different hit / false alarm ratios [cue validity]). Elderly and healthy, young adults completed the navigation task. As in real life, elderly walked slower and fell more often in pseudo-natural environments, suggesting the ecological validity of such safe experimental setups to test hazardous situations. Importantly, the vision aid reduced the risk of falling (by up to 33 %) for both age groups. Thus, our study highlights that a vision aid which augments reality could be a substantial contributor in preventing falls and related injuries, decreasing the financial burden on the health system and increasing life quality.

**Statement of relevance:** The present work tests the usefulness of a vision aid in decreasing fall risks. The study provides clear evidence that such vision aid substantially reduces incidences of obstacle collision thereby reducing the risk of falling and thus, related injuries. This effect appears to be age group independent, highlighting the relevance of the vision aid to function as a general medical tool for the larger public. Remarkably, the study also strongly suggests that real life behaviour can successfully be transferred to virtual environments, highlighting the potential of this method to investigate dangerous and potentially harmful situations in safe experimental environments.

## 1. Introduction

Injuries are not only painful but a major health risk and a financial burden on society. Present studies estimate 525.5 to 973 million injury-related incidents worldwide per year (Haagsma et al., 2016, 2020), rendering falling – the 2^nd^ most likely cause for unintentional injury deaths world-wide – roughly 4 % to 7 % of all injury-related incidents (WHO, 2021). The goal of the present study is to experimentally establish whether and how the cue validity of a visual augmented reality (AR)^1^ obstacle detection aid (i.e. the highlighting of hazardous objects in the environment) affects walking behaviour and thus, the risk of falling.

In 2021, 37.3 million falls were registered which required medical attention (WHO, 2021). Falls mainly result in a reduction of individual’s life quality, including hospitalisation, permanent disabilities, trauma induced anxiety and depression etc. (see e.g. Ang et al., 2020) and can even cause death (WHO, 2021: 684.000 fatal falls in 2021 with more incidents in the elderly population [> 60 years]). Falls also affect society in the form of increased needs for care giving, therapies and increased health system costs. For instance, the average costs per fall injury for people above 65 years were estimated with US $ 3611 and US $ 1049 in Finland and Australia, respectively. Although minor falls may be important for child development (i.e. improving balance), severe falls of children under 10 years produce costs of US $ 600 million each year (WHO, 2021). Thus, preventing falls can increase individual’s life quality, reduce long-term disabilities across different age groups and drastically reduce health care costs in general.

Below, we will focus on elderly people, as they have a higher likelihood of experiencing severe consequences after falling. Impaired visual, cognitive and muscle functions of elderly have frequently been linked to an increased risk of falling (de Boer et al., 2004; for review see Saftari & Kwon, 2018). For instance, elderly with lower binocular and lower distant visual acuity have a higher risk of multiple falls and suffering more severe injuries after falling (Klein et al., 2003; Koski et al., 1998; Lamoreux et al., 2008; Lord & Dayhew, 2001). Furthermore, poor vison was found to increase the relative risk of hip fractures (Felson et al., 1989). However, most of these studies are based on retrospective reports after the fall occurred. Thus, extrinsic fall risks – their influence on falls and their detection – are typically overlooked in fall prevention studies (Rajagopalan et al., 2017). Recent research started focussing on developing fall detection and fall prevention methods. Importantly, fall detection methods (Bian et al., 2015; Bourke et al., 2010) have major limitations: they only improve post-fall healthcare. In contrast, fall prevention methods use e.g. motion sensors to predict abnormal walking patterns (Howcroft et al., 2017; Majumder et al., 2013) or cameras and lasers mounted onto participant’s shoes to detect hazardous objects (Lin et al., 2017). Here, sounds are typically used to inform participants about a change in their walking pattern or the imminent collision with hazardous objects. However, an acoustic warning signal may increase alertness but does not necessarily facilitate the identification of the relevant object people have to bypass. Furthermore, participants have to be willing (and remember) to wear these extra devices. However, some devices are uncomfortable, result in bruises or are simply not user-friendly enough and thus, are sometimes avoided to wear in everyday life (Mathie et al., 2004).

To this end, acceptance and use of obstacle detection aids might be increased by extending devices frequently used in daily life, such as prescribed glasses^2^. Importantly, the incorporation of augmented reality into everyday devices – such as glasses – would require computer-aided detection of hazardous objects in our environment. As such algorithms can be computationally costly, we experimentally test how precise such detection algorithms have to be to reduce the risk of falling. To this end, participants guided an avatar (first person) through a virtual city and had to avoid hazardous objects. Objects were highlighted with different validities (e.g. “no highlighting” vs. “90 % correct highlighting and 0 % incorrect highlighting, below referred to as hits and false alarms, respectively). In general, we hypothesised that higher cue validity should result mainly in a reduction of collisions with hazardous objects (for more detailed hypotheses, see Analyses section below). Additionally, the augmented vision aid should compensate for physical disadvantages (e.g. decreased visual acuity); thus, elderly people should particularly benefit.

## 2. Methods

### 2.1. Participants

We collected data from two populations: young (18 to 35 years) and elderly (55 years and above). Based on a power of .99 and a small effect size (f = .1), we calculated a total sample size of 170 participants (85 per group) using G*Power 3 (Faul et al., 2007). Note that we set the sample goal for both groups to 90 participants to have fully counterbalanced datasets (see below). However, it was challenging to recruit young and elderly people during the pandemic. To compensate for this issue, we changed the entry year for the elderly from originally 65 to 55 years. However, this countermeasure did not fully solve the abovementioned problem and we were only able to measure roughly half of the intended sample in the elderly population during the funding period.

The young group served as a healthy control group. Participants were required to be free of colour blindness, walking impairment, hearing impairment, hand and motor skill impairment, motion sickness and mental or neurological disorders and had to have normal or corrected to normal vision and fluent German language skills. We could not collect data for one participant reporting a mental disorder while filling in the questionnaires. We collected and analysed data from 84 young participants (62 females, mean age 23.1 ± 3.98 SD; 12 left-handed).

For the elderly group the criteria had to be less strict, as this would otherwise have resulted in exclusion of most participants. Participants were required to have a minimum of 50 % vision (with visual aids) and high German language skills. Further, they were required to be able to understand speech (e.g. with the help of a hearing aid), walk for at least an hour (including short breaks) and be able to move and to grasp with both hands and arms (over a longer period of time, as they needed to use the joystick and mouse). They had to be free of colour blindness, motion sickness and psychological disorders. For 11 participants we could not collect data as they were unable to use the controls properly (7 people) or were not able to finish the short training levels in an appropriate time (4 people). We collected and analysed data from 40 elderly participants (20 females, mean age 68 ± 5.73 SD; 0 left-handed).

The study was approved by the ethics committee of the Otto-von-Guericke-University Magdeburg. It was carried out in careful compliance with the applicable Corona requirements. Depending on current government restrictions, 3G (vaccinated, recovered or tested) or 2G (vaccinated or recovered) rules applied (for elderly, we always used 2G restrictions to increase participant’s safety). In addition, all experimenters were tested for Covid-19 on the same day they measured participants. Participation was voluntary. Participants either received €8 per hour or student credits, based on their choice. During the experiment, all participants wore medical face masks when in contact with the experimenters. While completing the experiments, participants were sitting alone in a room with open windows and free to remove their mask.

### 2.2. Apparatus

We ran the experiment in two different laboratories (i.e. parallel measures) to be able to collect two instead of one dataset whenever multiple participants could only register for one particular time slot. In Laboratory 1, the experiment was run on an Acer ConceptD laptop (model number: CN517-71). For movement control, we used a Thrustmaster joystick (model number: T.16000M FCS) and a Microsoft Notebook Optical Mouse (model number: 1020). For sound, we used Creative speakers (model number: MF1695). In Laboratory 2, we used an AOC brand monitor (model number: 2436Vwa) in combination with a desktop computer. For movement control, we used again a Thrustmaster joystick (model number: T.16000M FCS) and a Dell mouse (model number: MOC5OU 0XN967). For sound, we used Logitech Z130 speakers (model number: 1-00098). Screens in both laboratories were set to a resolution of 1920 × 1080 pixels. As both screens differed in size (laptop: 31.1 × 21.5 cm versus monitor: 51.8 × 29.4 cm), we adjusted the visual angle (37.3 ° for the diagonals) across setups by placing participants in 65 cm versus 89 cm distance, respectively. The stereo loudspeakers’ volume was adjusted to 73.3 dB(A) in both laboratories using a 1 kHz sinusoidal test sound. We collected data of 117 participants (80 young, 35 elderly) in Laboratory 1 and of the remaining 9 participants (4 young, 5 elderly) in Laboratory 2.

Participants used the joystick for moving forward/backward and the mouse for changing the viewing direction. All participants operated the joystick with their left hand and the mouse with their right hand. The more the joystick was tilted forwards or backwards, the faster the avatar walked in either direction (see next section for maximum walking speed).

### 2.3. Stimuli, Avatar and visual obstacle detection aid

We used the Unity Engine (version 2019.4.16f1) by Unity Technologies for our study. Participants were required to walk through a virtual city with different districts (i.e. areas representing different distinct parts of a city, from suburban areas to financial districts; see Figure 1 for examples). We mainly used low to medium polygon assets to create a realistic looking city while keeping performance high (i.e. > 60 frames per second).

**Figure 1:**
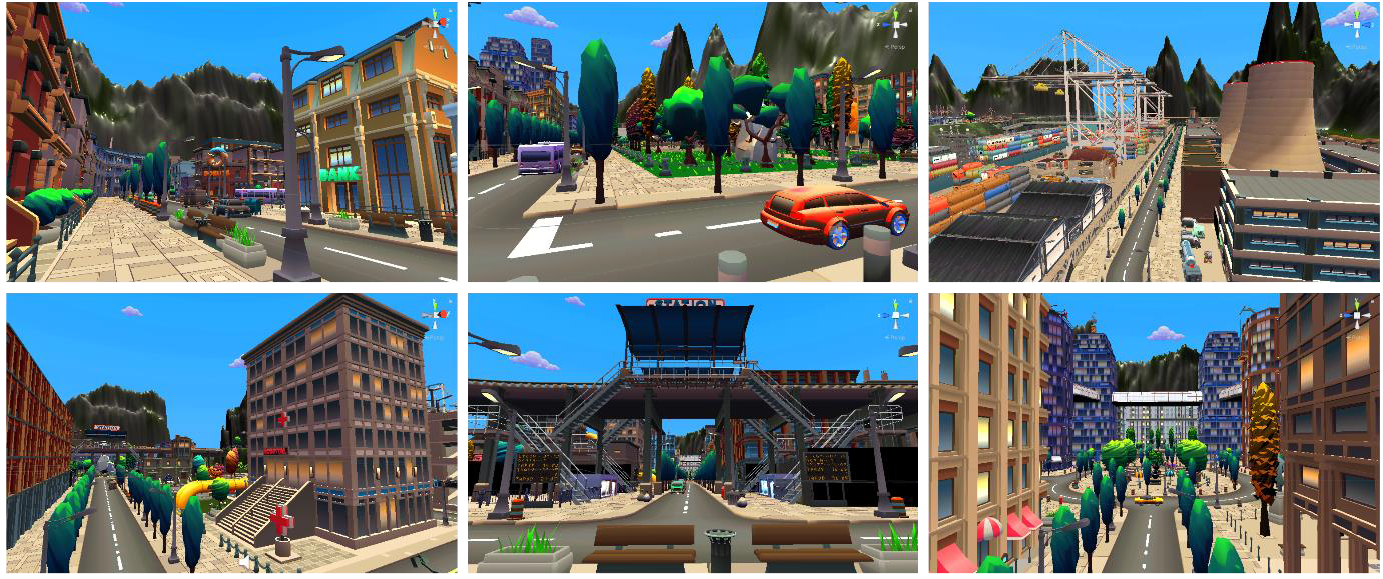
Examples with view from above for different city districts like harbour and financial district (right column).

To create the city, we used the Toon Desert, Toon Harbor Pack, Toon City, Toon Industries, Toon Nature Assets, Toon Gas Station assets from SICS Games and Polygon Fantasy Kingdom, Polygon Farm, Polygon Nature assets from Synty Studios. Boxophobic’s Polyverse Skies asset was used to create a dynamic sky box. The Cinemachine and Cinemachine Path assets were used to create moving trains and cars. To increase ecological validity, we presented car noises, ambient sounds (sounds of bustling cities, nature ambient sounds in parks, ship noises in the harbour area etc.) and foot step noises. To this end, we used the following assets: 2D Urban Cars (free sample) from Loonybits, 96 General Library (free sample pack) and 44.1 General Library (free sample pack) from InspectorJ Sound Effects, Engines by Kristian Grundström, Nature Sound FX by lumino, Pro Sound Collection by Gamemaster Audio and the Ultimate Sound FX Bundle 19 by Sidearm Studios. To have a visible player body (avatar), we used the The Wolf3D ReadyPlayerMe asset from the Unity Character Pack. Mixamo (https://www.mixamo.com/) was used for the walking animations of the avatar. The Internet sources of the assets are listed in Supplement 1. Additional objects and functionality (e.g. HoloLens simulation, movement, level transitions, stumbling and falling etc.) were custom designed and programmed.

Participants walked through the city in first person view (see Figure 2). The controlled avatar was fully animated and visible when looking down (to provide a more natural feeling to the experiment). In addition to walking animations, we added feedback for object collision in the form of stumble and fall animations. Fall animations included the fall as well as standing up again. The animations completed automatically and during that time, participants could not move the avatar (as in real life).

**Figure 2:**
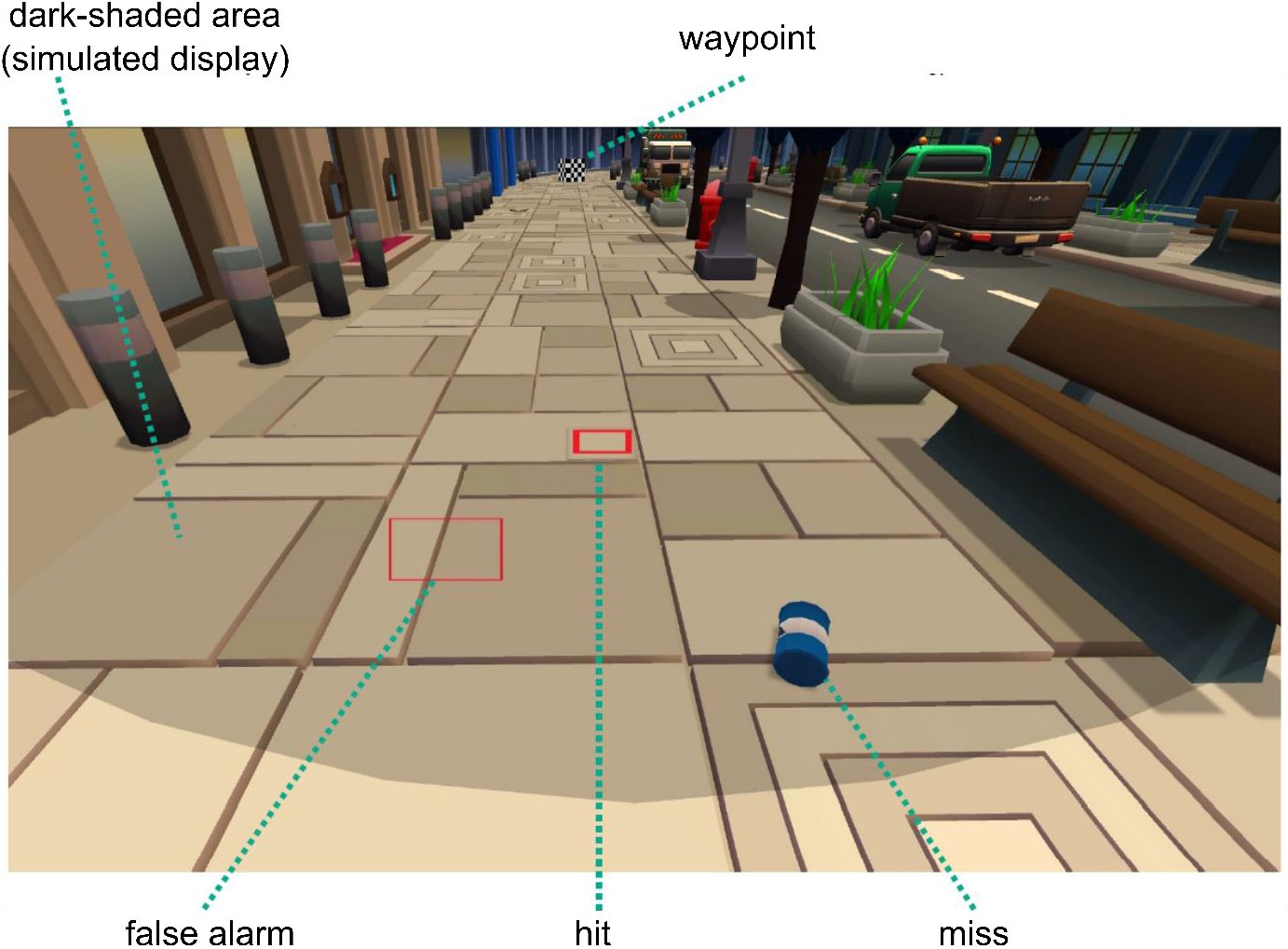
Example of a first-person view during the experiment. Participants always saw the visor/display of the vision aid (dark shaded area) and had to reach the next waypoint (top). In the “normal vision” condition nothing was highlighted. As soon as the vision aid was switched on, highlighting could be either correct (i.e. hits; here a barely visible stone plate not properly aligned with the side walk surface), incorrect (false alarms; false highlighting when no danger of falling is present) or missing (i.e. miss; whenever potential stumble objects were not highlighted).

The walking speed of the avatar was limited to a maximum of 3.57 Uunits/s (Unity units per second, ∼ 2.5 m/s) in accord with reports about average and maximum walking speeds (Bohannon, 1997). Note that one Uunit typically stands for 1 m in real life. However, the used assets to create the city were not properly scaled to Unity units. Based on the standard sizes of doors, fences, cars etc. we determined an approximated conversion factor of 1 Uunit = .7 m (1.43 Uunits = 1 m) for the assets used.

As in real life, participants used the side walk for navigating through the city. The side walk always consisted of three lanes (see Figure 2). Between two waypoints (check points participants had to pass, see below), we always placed a total of 15 “stumble objects”, with each lane containing three hard (more difficult to spot), one medium and one easy (more easily spotted) object (equals 5 stumble objects per lane). Stumble objects were manually and randomly placed between waypoints with the only restriction that participants were somehow able to continue walking (i.e. not all three lanes are blocked by obstacles at the same time). Stumble objects were e.g. stone plates, stones and twigs (see Figure 3 for examples). In addition, we used “decorative objects” to increase ecological validity and to avoid participants walking e.g. always on the curb. These decorative objects also triggered a collision but resulted only in a stumble animation. Decorative objects included benches, lanterns, traffic lights, rubbish bins, fences etc. (see Figure 2, right). While the stumble and decorative objects were used to simulate the hit rate of the visual aid, we placed empty objects (meaning objects that had no impact on collision and were not rendered, thus, invisible) to simulate false positive detections (see Figure 2). The distribution of empty objects in all lanes was identical to the stumble object ratios (3 hard, 1 medium and 1 easy object per lane between two waypoints).

**Figure 3:**
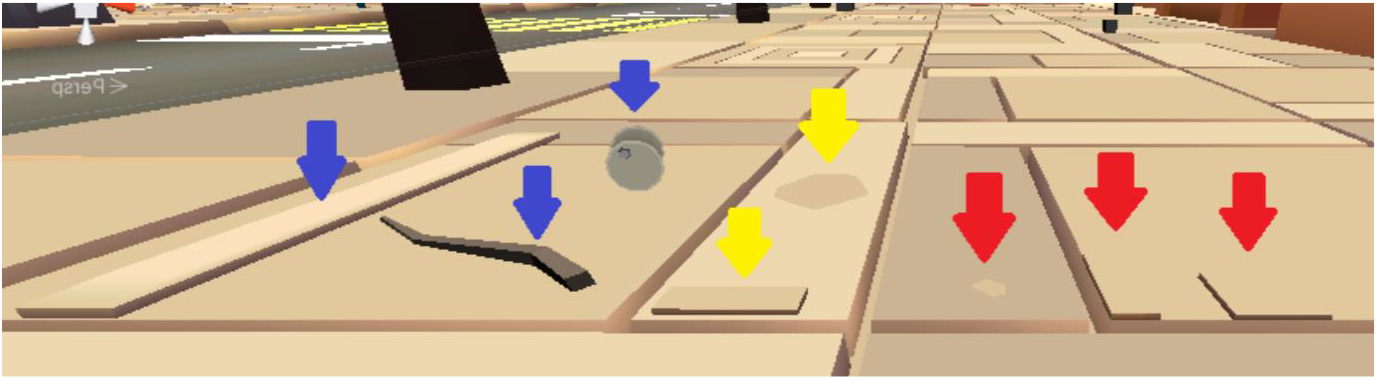
Example for easy- (blue arrow, left), medium- (yellow arrow, middle) and hard-to-detect (red arrow, right) stumble objects.

We created three custom designed objects: the waypoints (cubes with chequerboards pattern) participants had to pass, the visor of the simulated vision aid and the rectangles used for highlighting (see Figure 2). To render our results readily testable in real environments, participants always saw a visor as they would when wearing current state-of-the-art AR equipment, i.e. Microsoft’s HoloLens 2. Importantly, not only the visor but also the detection range of the vision aid was based on the current specifications of the HoloLens2 (currently, the farthest developed wireless AR device). Like the HoloLens2, our simulated obstacle detection aid had a detection radius of approx. 5 m in a 43° by 29° field of view (FoV was centred on the centre of the screen). Hence, not the whole screen (visual field) was used for detection of hazardous objects (based on current AR limitations). Note that with larger FoVs (as potentially implemented in future augmented reality devices) more objects could be detected in the environment; thus, by restricting ourselves to currently available devices, we likely did uncover the full potential current but not necessarily future augmented glasses, which likely have larger FoVs. For highlighting, we used red rectangles surrounding the potential hazardous objects to induce a pop out effect (see Figure 2) and most effectively manipulate attention to these objects (see e.g. Ball et al., 2014; Treisman & Gelade, 1980; Wolfe et al., 1989).

The cue validity of the obstacle detection (see Table 1) was manipulated by highlighting different percentages of the stumble and decorative objects (i.e. hits) and empty objects (i.e. false alarms). For instance, if the hit rate was set to 50 % and the false alarm rate 40 %, 50 % of the meaningful (could be collided with) stumble and decorative objects were highlighted and 40 % of the empty (non-existing) objects were highlighted whenever they were in the detection range of the obstacle detection aid. The chosen ratios of hits and false alarm rates for cue validity (see Table 1) were meant to sample, and thus simulate, a wide range of reasonable hit and false alarm rates (since it is not reasonable to assume that any computational detection algorithm always produces false alarm rates of 0 %) as well as a wide range of their possible combinations within the ROC area of the visual aid (i.e. its detection sensitivity).

**Table 1:** Different cue validities for obstacle detection in the present study, ranked by their d′ value (discrimination sensitivity index in signal detection theory) from lowest (1) to highest (9) validity. Note that the combinations 0:0 and 90:90 actually produce the same d′ value (d’ = 0). However, we ranked the 90:90 condition higher as 90 % of the hazardous objects were highlighted in this condition (instead of no highlighting).

| Cue validities | 1 | 2 | 3 | 4 | 5 | 6 | 7 | 8 | 9 |
| --- | --- | --- | --- | --- | --- | --- | --- | --- | --- |
| *hit rate (%)* | 50 | 0 | 90 | 60 | 30 | 50 | 80 | 70 | 90 |
| *false alarm rate (%)* | 90 | 0 | 90 | 40 | 10 | 20 | 40 | 20 | 0 |
| *resulting $d'$ of vision aid* | -1.28 | 0 | 0 | 0.51 | 0.76 | 0.84 | 1.1 | 1.37 | 7.28 |

### 2.4. Questionnaires

In this study, we focused on the effects of the vision aid on walking behaviour and whether these effects differ between young and elderly people. To account for any age-related influences on the collision data, we conducted a general health questionnaire and test battery following suggestions by de Boer et al. (2004) and a post-hoc questionnaire. The goal was to assess potentially meaningful covariates and thus, to be able to differentiate between their influences and our effect of interest (for an overview, see Table ***2***).

**Table 2:** List of all covariates and their potential influence on the data.

| Covariate | Reasoning |
| --- | --- |
| *visual acuity* | Participants with lower visual acuity might not be able to properly see the warning cues and objects in the surroundings and might thus, stumble more often than others and/or walk slower, independently of the effects of the vision aid. |
| *colour vision* | Participants with lower colour vision scores might not be able to properly see the warning cues. |
| *vision aid usage* | We combined the scores of “how much the vision aid information helped” and “whether participants relied on it” as they were highly correlated. Higher values indicate that participants relied more on the HoloLens and that it was more helpful. Participants with lower scores might subjectively be less inclined to use the information the vision aid provides. |
| *ignored objects* | Participants might be inclined to forget about monitoring hazardous objects which are not highlighted. |
| *physical fitness* | We combined the scores of “physical fitness self-evaluation” and “LAPAQ” as they were highly correlated. Lower values indicate less fitness and mobility. Participants with lower scores might have irregular and slower walking behaviour as they are less used to physical activities. |
| *cognitive fitness* | Lower cognitive fitness, as measured by the MMSE, might affect attention to relevant objects and thus, walking behaviour |

**Table 3:** Overview for results with covariates (right) for all analysed variables and effects of interest. Note that we list the important numbers for effects without covariates on the left. All effects including factor cue validity (i.e. main effect and interaction) were GG corrected.

|  |  | Without covariates |  |  | With covariates |  |  |  |  |
| --- | --- | --- | --- | --- | --- | --- | --- | --- | --- |
| | | F | p | $\eta_p^2$ | F | df | p | $\eta_p^2$ | BF<sub>incl</sub> |
| **Collision** | *cue validity* | 13.561 | < .001 | .1 | 13.977 | 6.55,759.93 | < .001 | .108 | 3.249E+15 |
|  | *Age* | 161.666 | < .001 | .57 | 72.342 | 1,116 | < .001 | .384 | 1.078E+13 |
|  | *Interaction* | .996 | .43 | .008 | 1.159 | 6.56,799.66 | .325 | .01 | 0.007 |
| **walking speed** | *cue validity* | 1.917 | .069 | .015 | 1.786 | 6.539,758.556 | 0.092 | 0.015 | 3.6*10<sup>-1</sup> |
|  | *Age* | 79.656 | < .001 | .395 | 35.715 | 1, 116 | < .001 | 0.235 | 7.7*10<sup>7</sup> |
|  | *Interaction* | .679 | .681 | .006 | 0.27 | 6.539,758.556 | 0.96 | 0.002 | 5.9*10<sup>-1</sup> |
| **Distance** | *cue validity* | 1.51 | .210 | .012 | 1.107 | 3.094, 358.93 | 0.347 | 0.009 | 3*10<sup>-3</sup> |
|  | *Age* | 3.086 | .081 | .025 | 0.122 | 1, 116 | 0.728 | 0.001 | 3.6*10<sup>-1</sup> |
|  | *Interaction* | .771 | .515 | .006 | 0.328 | 3.094, 358.93 | 0.811 | 0.003 | 4*10<sup>-3</sup> |

The questionnaire/test battery included tests for vision acuity and colour vision (to this end, we used two freely available online tests [run on the same monitors used for the experiment]: https://www.brillen-sehhilfen.de/sehtest/, https://www.sehtestbilder.de/online-sehtest-kostenlos.php), a self-evaluation of their physical fitness (difficulties with daily activities), the LASA Physical Activity Questionnaire (LAPAQ; indicating their mobility), and the Mini-Mental State Examination (MMSE) to assess cognitive impairments.

The post-hoc questionnaire assessed participants’ subjective experience, i.e. to which extent (in percent) the visual aid was helping them (0 % = no help at all, 100 % = absolutely helpful), to which extent they relied on the visual aid (five-point Likert scale) and to which extent they ignored dangerous objects outside of the highlighted areas (five-point Likert scale).

### 2.5. Procedure

The experiment consisted of 6 stages. First, participants provided written informed consent. Second, motion sickness was tested for by letting participants move freely through the first training level, asking them to stumble and fall over objects and to report their subjective states afterwards. Third, participants completed the general health questionnaire and test battery (see section 2.2.). If participants got motion sick or the data of the test battery indicated that participants did not meet the participation criteria, we stopped the experiment.

Fourth, participants completed a training to familiarize themselves with the controls, potential collision objects and the visual aid. To this end, we created two levels which were used only for training. In the first training level nothing was highlighted (= normal vision). In the second training level, we always used a hit rate of 90 % and a false alarm rate of 0 %. In case participants from the young group required more training, the two levels were repeated. Given that elderly people have typically less contact with computers, both training levels were repeated up to a maximum of four times. Although elderly people might be generally slower in learning the controls, as they have less practise with computers, we needed to assure that they complete the experiment in an acceptable amount of time. To this end, we set the criterion of ∼ 8 min per level in the last training run (typically resulting in 5h long sessions including breaks and questionnaires). In case, elderly participants took significantly longer than the criterion (>= 10 min per level), we stopped the experiment and the participant was excluded from the experiment.

Fifth, participants completed a total of 18 levels in the main experiment. Again, they were asked to walk through each level as quickly and safely as possible, i.e. without falling. The main focus was on the aspect of safety without neglecting speed. Participants did not navigate randomly but had to walk through waypoints (rotating cubes with a chequerboards pattern). If one waypoint was “collected” (i.e. hit by the avatar), the next one appeared and the current one disappeared. After collecting the last waypoint, the avatar transitioned to the next level. If the participants hit one of the obstacles (see point 2.3.), the avatar stumbled or fell. For every 5 stumbles the character fell one time. We used this ratio based on piloting data as more fall animations resulted in a drastic increase of motion sickness, due to their duration and shakiness. By using less fall animations and more stumble animations, we were able to reduce motion sickness and thus, collect more datasets while still providing meaningful feedback for collisions with objects in the environment.

Participants could activate a “safe walk” mode (25 % of the current walking speed) by pressing the left mouse button. We included this option for situations in which people – like in real life – slow down to pass obstacles safely (e.g. navigating smaller abrupt height distances in terrain). For the duration of the button press they were able to walk over obstacles without triggering stumbles or falls. However, their progression was slowed down significantly, motivating them to only use this option when necessary to adhere to the speed-criterion. Alternatively, they could always navigate around the obstacles. In case they had to cross the street to reach the next waypoint, they were always required to use the safe walk mode for the transition from side walk to street and vice versa. On the street itself, they could walk normally. If they missed to activate the safe walk mode during transition, they stumbled. Importantly, participants were instructed that cars will not stop for them and they were required to monitor car movements (as in real life) before crossing the street. If a car crash happened, the avatar respawned at the last waypoint after a five second penalty.

To test the effect of AR glasses on walking behaviour, we manipulated the cue validity of the obstacle detection across levels. We simulated nine different cue validities (i.e. specific combination of hit and false alarm rate; see Table 1). Validities always changed after two levels. The change of validity was announced in form of a transition screen. However, participants never received information about the exact cue validity after change, as such feedback would also be impossible in real life. The sequence of the nine cue validities was counterbalanced across participants using the Latin square design method (Bradley, 1958), resulting in 18 possible sequences. Note that the sequence of the levels themselves was always fixed (e.g. harbour followed by finance district followed by rural area etc.). However, the variable of interest – the obstacle detection cue validity – was counterbalanced for each level.

After completing all levels, participants answered the post-hoc questionnaire. The experiment took approximately 2 to 3 hours for participants in the young group and 3.5 to 5 hours for participants in the elderly group.

We provide two example videos of the experiment in the Supplement.

### 2.6. Analyses

Data was processed and analysed using the MATLAB 2017b software (The MathWorks Inc.) and JASP software (JASP Team, version 0.14.1). Whenever required, ANOVA results were Greenhouse-Geisser (p_GG_) corrected. We also used JASP to calculate the effect size (η_p_ ^2^), conduct post-hoc tests (Bonferroni corrected; p_Bonf_), and to calculate the Bayes factors (matched model effects; standard settings for priors and samples: r scale fixed effects = .5, r scale random effects = 1, r scale covariates = .354, samples = “auto”) in favour of the alternative hypothesis (BF_incl_). Note that BF_incl_ smaller than 1 are (more) in favour of the null hypothesis (Wagenmakers et al., 2018).

We used three different measures to analyse participants walking behaviour (for explanation see below): *collision frequency, walking speed*, and *relative distance*. Note that we sorted conditions (i.e. cue validities) for our analysis according to Table 1. As each condition was used in two consecutive levels, we averaged data across levels.

First, we calculated the *collision frequency*. To this end, we normalised the number of falls/stumbles with the walked distance (walked Uunits) in the respective level. Thus, higher values indicate more collisions. Note that we analysed collision frequency instead of the raw number of collisions due to different level and thus, walking distances (e.g. 5 collisions on 5 units walked would have the same frequency as 20 collisions on 20 units walked although the total amount of collisions would appear to differ). *Specific hypotheses:* If the vision aid is effectively manipulating walking behaviour, participants should take the warning cues into account and in turn, the frequency of collisions with hazardous objects should be reduced as compared to the no-highlight condition. If the specific cue validity is of importance, collision frequency should decrease with higher cue validity.

Second, we calculated the *walking speed* (average Uunits/s) with higher numbers indicating faster walking speed. Note that we calculated the average walking speed by excluding frames with animations (e.g. stumbling or falling); otherwise walking speed would be artificially decreased by and correlated with events during which participants were actually not walking (i.e. collisions). *Specific hypotheses:* Participants should walk faster under ‘highlighting’ conditions than under ‘no highlighting’ conditions. If higher cue validities actually reduce the frequency of collisions, participants should also be more confident and thus, walk faster under these conditions. Furthermore, if our walking simulator successfully mimics reality, elderly should walk slower than young people given their natural difference in walking speed.

Third, we calculated the *relative distance*. To this end, we normalised the true distance walked by the optimal level distance (i.e. a straight line between waypoints; average optimal level distance was 333.2 Uunits ± 31.7 Uunits SD) to account for general differences in level distances. Thus, higher values indicate longer distances walked. *Specific hypotheses:* Highlighting might result in higher relative distances, as people navigate around potentially hazardous objects instead of walking in a straight line.

To test our hypotheses, we ran separate repeated-measures ANOVAs for each variable of interest with within-subject factor *cue validity* (1-9), between-subject factor *age group* (young, elderly), once without and once with the added covariates *visual acuity, colour vision, vision aid usage, ignored objects, physical fitness and cognitive fitness* (for an overview, see Table ***2***) to account for potential effects of age-related burdens, driving or obscuring the effects of interest. Note that we report both versions of the analyses to show that the effects are analysis-independent and robust.

## 3. Results

The analysed data are available in the supplementary file.

### 3.1. Without covariates

As hypothesised, higher cue validities reduced the frequency of collisions with hazardous objects (F(6.56, 799.66) = 13.561, p_GG_ < .001, η_p_^2^ = .1, BF_incl_ = 3.2*10^15^). Collisions per 100 Uunits were reduced from 2 collisions (no vision aid) to 1.4 collisions (highest cue validity). This is a reduction of collision frequency by 33 %. Further note that hit rates larger than 70 % typically resulted in a reduction of collision frequency. More importantly, hit rates of 90 % improved performance even when paired with a false alarm rate of 90 % (cue validity 3) and were not significantly different from the highest cue validity (cue validity 9). As in real life, elderly participants collided more frequently with objects in the surrounding (stumble and decorative) than young participants (average per 100 Uunits: elderly – 3.2 collisions, young – 1.1 collision; F(1,122) = 161.666, p < .001, η_p_^2^ = .57, BF_incl_ = 6.2*10^20^). Remarkably, the interaction of factors cue validity and age group was non-significant (F(6.56,799.66) = .996, p_GG_ = .43, η_p_^2^ = .008, BF_incl_ = .007), highlighting the usefulness of the vision aid for young and elderly. The data plots for significant effects and post-hoc comparisons are shown in Figure 4:

**Figure 4:**
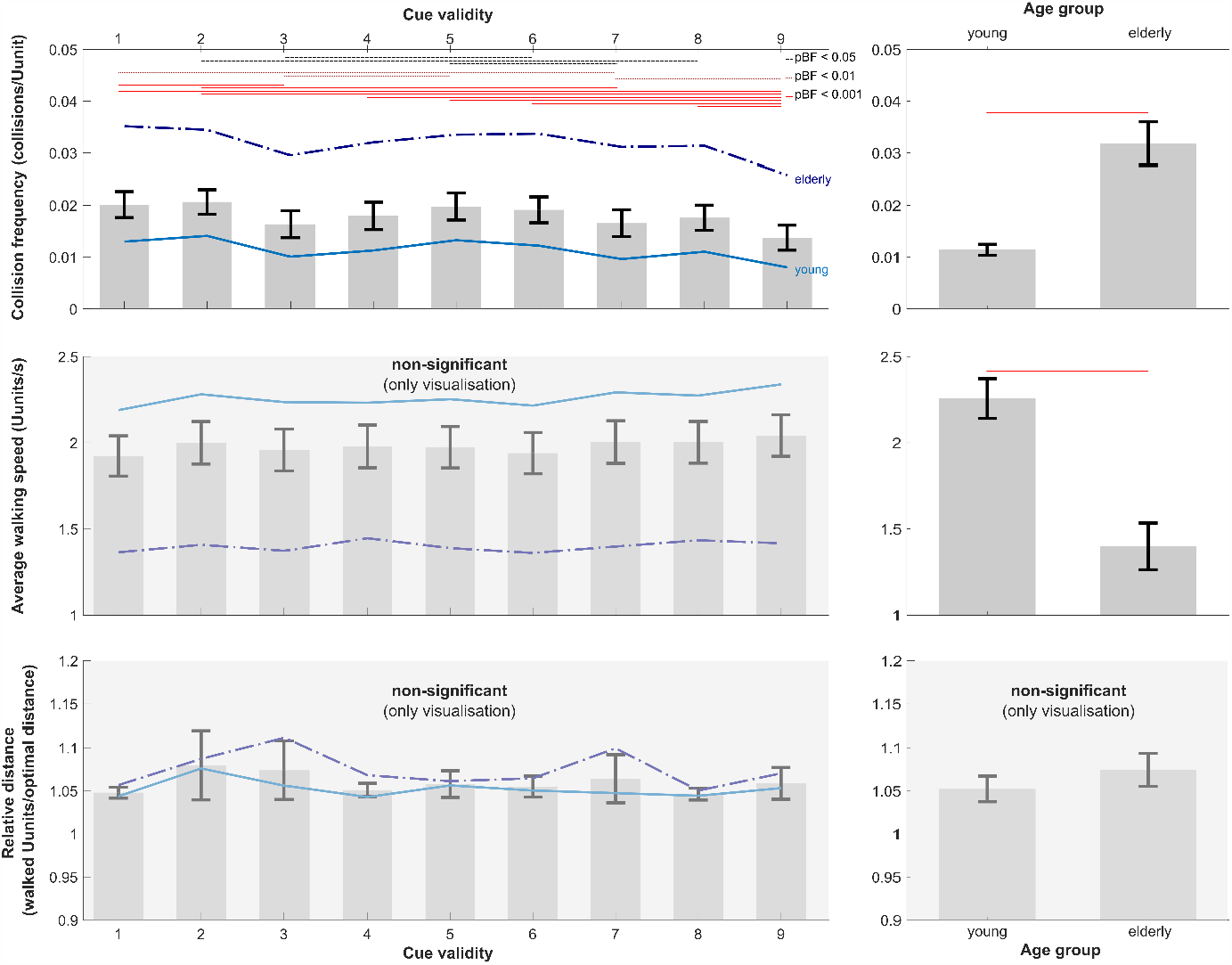
Depiction of behavioural results (top: collision frequency, middle: walking speed, bottom: walked distance). Note that we also plotted the data for non-significant effects. Left bar graphs show the effect for the factor level of cue validity and the right bar graphs for factor age group. Bars represent group mean averages (with error bars indicating 95 % confidence intervals). The only significant effect of cue validity was found for the collision frequency data: here we also depict significant post hoc comparisons in form of bars on top of the graph. Bonferroni corrected post hoc p-values are colour coded as follows: bright red + solid: p < 0.001, dark red + dotted: p < 0.01; black + dashed: p < 0.05. For illustration purposes, we also present the average values of each age group (elderly: dark blue + dashed/dotted; young: bright blue + solid) to highlight their similarity resulting in the absence of interaction effects.

For *walking speed*, we only found a trend indicating that higher cue validities may foster increased walking speeds (F(6.56,800.71) = 1.917, p_GG_ = .069, η_p_ ^2^ = .015, BF_incl_ = .364). Most importantly, the difference between young and elderly was once more significant and elderly participants walked – as hypothesised – on average slower than young participants (F(1,122) = 79.656, p < .001, η_p_ ^2^ = .395, BF_incl_ = 5.7*10^11^). The interaction of factors cue validity and age group was again non-significant (F(6.56,800.71) = .679, p_GG_ = .681, η_p_ ^2^ = .006, BF_incl_ = .004). As for the collision data, we present the two main effects in Figure 4: .

For *relative walked distance* all results were non-significant (F(3.1,378.14) = 1.51, p_GG_ = .21, η ^2^ = .012, BF_incl_ = .003; trend for *age group:* F(1,122) = 3.081, p = .081, η_p_ ^2^ = .025, BF_incl_ = .712; *interaction:* F(3.1,378.14) = .771, p_GG_ = .515, η_p_ ^2^ = .006, BF_incl_ = .004). As for the collision data, we present the two main effects in Figure 4: for illustrative purposes.

### 3.2. With covariates

The results for the effects of interest with covariates are listed in ***Table 1*** next to the results without covariates. Note that all results are virtually identical when accounting for potential covariates. Thus, our results are independent of the type of analysis. For interested readers, we list all results for covariates in Supplement 2, as we were not interested in the covariates themselves.

## 4. Discussion

The present study demonstrates that augmented reality vision aids can reduce the risks of falling due to collisions with hazardous objects across age groups, rendering the vision aid a universal age-independent help in preventing falls.

For experimental testing purposes, we likely presented more opportunities to stumble than in real life. Nevertheless, our data suggests an approximate average reduction of fall frequency and total falls of 33 % across age groups (10 vs. 6 falls for young people and 25 vs. 19 falls for elderly people). Even if we assume that this number is significantly lower in real life scenarios, a reduction of falls by e.g. only 5 % would still have a significant impact. Reducing falls by a ‘mere’ 5 % would imply that roughly 2 million people wouldn’t have suffered fall related injuries in 2021, thus, increasing their life quality by avoiding post-injury health complications and psychological stress disorders. With an average cost of roughly US $ 2000 per injury case, US $ 4 billion could have been saved. These numbers highlight the possible implications and benefits of implementing an augmented reality vision aid in daily life.

Another important finding of this study is that participants’ behaviour in our virtual reality appears to relate to real life behaviour. Elderly people walked with approximately 3-3.5 km/h and younger people with 5-5.5 km/h in our simulation. These numbers correspond to slow and fast walking speeds one would expect for elderly and young people in real life (Bohannon, 1997; Bohannon & Andrews, 2011; Kyrdalen et al., 2019; Studenski et al., 2011). Although not of direct interest to the research question at hand, we also found that the number of collisions generally increased e.g. with impaired vision and lower cognitive fitness, factors typically associated with age and typically correlating with risks of falling (de Boer et al., 2004; for review see Saftari & Kwon, 2018). In line, elderly people also showed higher rates of collisions, independent of whether covariates were accounted for. Thus, these findings also mirror real world data (Talbot et al., 2005). Given the overall similarities across age groups (i.e. no interaction effect) and the similarities in behaviour between simulation and real life, we draw two conclusions. First, it is unlikely that differences in technical affinity (familiarity in working with technical devices) across age groups drove our results. If such differences had had an impact on behaviour, we would have expected some form of interaction effect which was absent. While we cannot completely rule out that differences in technical prowess might have had partially decreased speed and increased collisions in the elderly group, our procedure (assuring that elderly people are able to use the mouse and joystick properly) and the close relation to real life data renders technical prowess rather unlikely to be the origin of our results. Second and most importantly, our results suggest that virtual environments appear to be suitable to test more hazardous scenarios (such as increased fall risks) in safe experimental settings instead of running the same test directly under less safe real-life conditions.

While these results are promising, tests of a real augmented reality vision aid still need to be conducted. For instance, we currently do not know whether a vision aid could affect walking speed. While our results suggest that walking speed is unaffected by augmented reality information, it is possible that people felt too save in the simulation. This might have resulted in participants walking with their average maximum speed irrespective of cue validity. Whether this is true can only be tested by transferring the experiment to real life. However, another reason for this result might be that we changed cue validity always after two level. Thus, there was no continuity and people had to adapt to new settings repeatedly. Following this reasoning, differences in walking speed might arise when only two conditions are presented and compared, i.e. no highlighting versus one helpful cue validity, providing participants with continuous, identical feedback, thereby potentially increasing the effect of walking speed differences.

Furthermore, it is currently unclear what happens when people stop using the vision aid. One possibility is that the aid improves obstacle detection in the long term and thus, reduces falls even when turned off. However, the opposite might be true. After becoming more confident in their walking pattern during training with the vision aid, the missing obstacle detection might now result in more collisions than before training. Future studies are required to test these hypotheses.

Finally, one might ask themselves why a new device is necessary. As outlined in the introduction, most current devices only detect falls after they have happened and ‘the damage is already done’ (Bian et al., 2015; Bourke et al., 2010; Rajagopalan et al., 2017). Other devices detect hazardous objects but are mainly meant for blind people (Dionisi et al., 2012; Laubhan et al., 2016; Nishajith et al., 2018; Simôes & De Lucena, 2016) and thus, provide no visual-spatial feedback (i.e. only spatially uninformative sounds or path descriptions), rendering the identification of objects a challenging endeavour. Additionally, people might not be willing to wear external devices for several reasons. For instance, mounting a camera or laser on ones shoes (Lin et al., 2017) might not be easily possible for elderly physically. Furthermore, it ‘might also look silly’ to wear such shoes, rendering elderly people less prone to use such devices continuously. In line, people appear less likely to wear existing devices which are uncomfortable or not user-friendly (Mathie et al., 2004).

By simply modifying existing glasses, which a large part of the elderly population has to wear standardly (Otte et al., 2018), and transforming them into a vision aid, which can be turned on and off with a simple button press, the aforementioned challenges in technical acceptance and usage would be easily circumvented. Given the pattern of results, the most relevant aspect of the detection algorithm appears to be the hit rate as even the condition with 90 % hit rate and 90 % false alarm rate improved performance. Thus, even if a multitude of objects (be they real obstacles or just false alarms) is highlighted, fall risk is reduced because the relevant objects are part of the highlighting and capture attention. Most importantly, on average, participants stated that they neither relied nor did not rely on the vision aid (neutral score of 3.1 ± .8 on five-point Likert scale) and that the vision aid only partially helped them (38.9 ± 26.8 %). Thus, our data suggest that even if people might be put off by multiple objects being highlighted and do not necessarily think that such vision aid helps them, we still see a reduction in obstacle collisions, rendering a visual obstacle detection aid a substantial health improvement.

## Supporting information

Video1_Example

ideo2_Example

## Acknowledgements

We thank Prof. Theodore Zanto for discussions inspiring this experiment.

## Declarations

### Author’s contribution (CRediT author statement)

**Felix Ball:** Conceptualization, Data curation, Formal analysis, Funding acquisition, Investigation, Methodology, Project administration, Resources, Software, Supervision, Visualization, Writing – original draft, Writing – review & editing **Judith Bomba:** Investigation, Software, Writing - Original Draft, Writing - Review & Editing **Annika Nentwich:** Investigation, Software, Writing - Original Draft, Writing - Review & Editing **Fabian Meier:** Investigation, Software, Writing - Original Draft, Writing - Review & Editing **Peter Vavra:** Conceptualization, Writing - Original Draft, Writing - Review & Editing **Toemme Noesselt:** Conceptualization, Supervision, Writing - Original Draft, Writing - Review & Editing

### Ethics approval

This study was approved by the local ethics committee of the Otto-von-Guericke-University, Magdeburg.

### Consent to participate and for publication

Informed consent was obtained from all individual participants included in the study.

### Open Practices Statement

The data and materials are available in the supplementary information files and upon request (e.g. codes). None of the experiments were preregistered.

### Data availability statement

The authors declare that the analysed data as well as potential supplementary analyses (not reported in the main manuscript) are available in the supplementary information files.

### Code availability

Data were analysed with JASP (freely available) and Matlab 2017b (Mathworks Inc.). Any relevant code is available upon request.

## Footnotes

1 Observers see computer-generated items in the environment (e.g. a web browser floating in the air) by means of an augmented reality device (e.g. glasses with an attached mini-computer projecting onto the glasses).

2 The following numbers highlight the general use of glasses in our society: In the US, roughly 25-30% of children (< 17 years) and over 65% of adults wear prescribed vision aids (The Vision Council, 2021; Villarroel, 2021). In Western European countries more than 50-60% of the adult population uses prescribed glasses (Michas, 2022). In addition, augmented reality technology is on the rise (Alsop, 2022), with a current estimate of 1 billion users (expected to increase to 1.73 billion users in 2024). Creations like SMART glasses (e.g. Google glasses), AR devices in the format of standard prescription glasses, make it highly likely that prescription glasses can be AR modified in the near future.

## References

Alsop, T. (2022). Global mobile augmented reality (AR) users 2024 | Statista. https://www.statista.com/statistics/1098630/global-mobile-augmented-reality-ar-users/

Ang, G. C., Low, S. L., & How, C. H. (2020). Approach to falls among the elderly in the community. Singapore Medical Journal, 61(3), 116–121. 10.11622/SMEDJ.2020029

Ball, F., Elzemann, A., & Busch, N. A. (2014). The scene and the unseen: Manipulating photographs for experiments on change blindness and scene memory Image manipulation for change blindness. BEHAVIOR RESEARCH METHODS, 46(3), 689–701. 10.3758/s13428-013-0414-2

Bian, Z.-P., Hou, J., Chau, L.-P., & Magnenat-Thalmann, N. (2015). Fall Detection Based on Body Part Tracking Using a Depth Camera. IEEE Journal of Biomedical and Health Informatics, 19(2), 430–439. 10.1109/JBHI.2014.2319372

Bohannon, R. W. (1997). Comfortable and maximum walking speed of adults aged 20-79 years: reference values and determinants. Age and Ageing, 26(1), 15–19. 10.1093/AGEING/26.1.15

Bohannon, R. W., & Andrews, A. W. (2011). Normal walking speed: a descriptive meta-analysis. Physiotherapy, 97(3), 182–189. 10.1016/j.physio.2010.12.004

Bourke, A. K., van de Ven, P., Gamble, M., O’Connor, R., Murphy, K., Bogan, E., McQuade, E., Finucane, P., ÓLaighin, G., & Nelson, J. (2010). Evaluation of waist-mounted tri-axial accelerometer based fall-detection algorithms during scripted and continuous unscripted activities. Journal of Biomechanics, 43(15), 3051–3057. 10.1016/J.JBIOMECH.2010.07.005

Bradley, J. V. (1958). Complete Counterbalancing of Immediate Sequential Effects in a Latin Square Design. Journal of the American Statistical Association, 53(282), 525. 10.2307/2281872

de Boer, M. R., Pluijm, S. M., Lips, P., Moll, A. C., Völker-Dieben, H. J., Deeg, D. J., & van Rens, G. H. (2004). Different Aspects of Visual Impairment as Risk Factors for Falls and Fractures in Older Men and Women. Journal of Bone and Mineral Research, 19(9), 1539–1547. 10.1359/JBMR.040504

Dionisi, A., Sardini, E., & Serpelloni, M. (2012). Wearable object detection system for the blind. 2012 IEEE International Instrumentation and Measurement Technology Conference Proceedings, 1255–1258. 10.1109/I2MTC.2012.6229180

Faul, F., Erdfelder, E., Lang, A. G., & Buchner, A. (2007). G*Power 3: A flexible statistical power analysis program for the social, behavioral, and biomedical sciences. Behavior Research Methods 2007 39:2, 39(2), 175–191. 10.3758/BF03193146

Felson, D. T., Anderson, J. J., Hannan, M. T., Milton, R. C., Wilson, P. W., & Kiel, D. P. (1989). Impaired vision and hip fracture. The Framingham Study. Journal of the American Geriatrics Society, 37(6), 495–500. http://www.ncbi.nlm.nih.gov/pubmed/2715555

Haagsma, J. A., Graetz, N., Bolliger, I., Naghavi, M., Higashi, H., Mullany, E. C., Abera, S. F., Abraham, J. P., Adofo, K., Alsharif, U., Ameh, E. A., Ammar, W., Antonio, C. A. T., Barrero, L. H., Bekele, T., Bose, D., Brazinova, A., Catalá-López, F., Dandona, L., … Vos, T. (2016). The global burden of injury: incidence, mortality, disability-adjusted life years and time trends from the Global Burden of Disease study 2013. Injury Prevention, 22(1), 3–18. 10.1136/INJURYPREV-2015-041616

Haagsma, J. A., James, S. L., Castle, C. D., Dingels, Z. V., Fox, J. T., Hamilton, E. B., Liu, Z., Lucchesi, L. R., Roberts, N. L. S., Sylte, D. O., Adebayo, O. M., Ahmadi, A., Ahmed, M. B., Aichour, M. T. E., Alahdab, F., Alghnam, S. A., Aljunid, S. M., Al-Raddadi, R. M., Alsharif, U., … Vos, T. (2020). Burden of injury along the development spectrum: associations between the Socio-demographic Index and disability-adjusted life year estimates from the Global Burden of Disease Study 2017. Injury Prevention : Journal of the International Society for Child and Adolescent Injury Prevention, 26(Supp 1). 10.1136/INJURYPREV-2019-043296

Howcroft, J., Kofman, J., & Lemaire, E. D. (2017). Prospective Fall-Risk Prediction Models for Older Adults Based on Wearable Sensors. IEEE Transactions on Neural Systems and Rehabilitation Engineering, 25(10), 1812–1820. 10.1109/TNSRE.2017.2687100

Klein, B. E. ., Moss, S. E., Klein, R., Lee, K. E., & Cruickshanks, K. J. (2003). Associations of visual function with physical outcomes and limitations 5 years later in an older population. Ophthalmology, 110(4), 644–650. 10.1016/S0161-6420(02)01935-8

Koski, K., Luukinen, H., Laippala, P., & Kivelä, S.-L. (1998). Risk Factors for Major Injurious Falls among the Home-Dwelling Elderly by Functional Abilities. Gerontology, 44(4), 232–238. 10.1159/000022017

Kyrdalen, I. L., Thingstad, P., Sandvik, L., & Ormstad, H. (2019). Associations between gait speed and well-known fall risk factors among community-dwelling older adults. Physiotherapy Research International, 24(1), e1743. 10.1002/pri.1743

Lamoreux, E. L., Chong, E., Wang, J. J., Saw, S. M., Aung, T., Mitchell, P., Wong, T. Y., & Wong, T. Y. (2008). Visual Impairment, Causes of Vision Loss, and Falls: The Singapore Malay Eye Study. Investigative Opthalmology & Visual Science, 49(2), 528. 10.1167/iovs.07-1036

Laubhan, K., Trent, M., Root, B., Abdelgawad, A., & Yelamarthi, K. (2016). A wearable portable electronic travel aid for blind. 2016 International Conference on Electrical, Electronics, and Optimization Techniques (ICEEOT), 1999–2003. 10.1109/ICEEOT.2016.7755039

Lin, T.-H., Yang, C.-Y., & Shih, W.-P. (2017). Fall Prevention Shoes Using Camera-Based Line-Laser Obstacle Detection System. Journal of Healthcare Engineering, 2017, 8264071. 10.1155/2017/8264071

Lord, S. R., & Dayhew, J. (2001). Visual risk factors for falls in older people. Journal of the American Geriatrics Society, 49(5), 508–515. http://www.ncbi.nlm.nih.gov/pubmed/11380741

Majumder, A. J. A., Zerin, I., Uddin, M., Ahamed, S. I., & Smith, R. O. (2013). smartPrediction. Proceedings of the 2013 Research in Adaptive and Convergent Systems on - RACS’ 13, 434–439. 10.1145/2513228.2513267

Mathie, M. J., Coster, A. C. F., Lovell, N. H., Celler, B. G., Lord, S. R., & Tiedemann, A. (2004). A pilot study of long-term monitoring of human movements in the home using accelerometry. Journal of Telemedicine and Telecare, 10(3), 144–151. 10.1258/135763304323070788

Michas, F. (2022). Spectacle wearers in Europe in 2020 | Statista. https://www.statista.com/statistics/711514/individuals-who-wear-spectacles-in-selected-european-countries/

Nishajith, A., Nivedha, J., Nair, S. S., & Shaffi, J. M. (2018). Smart cap-wearable visual guidance system for blind. 2018 International Conference on Inventive Research in Computing Applications (ICIRCA), 275–278. 10.1109/ICIRCA.2018.8597327

Otte, B., Woodward, M. A., Ehrlich, J. R., & Stagg, B. C. (2018). Self-Reported eyeglass use by US Medicare beneficiaries aged 65 years or older. JAMA Ophthalmology, 136(9), 1047–1050. 10.1001/jamaophthalmol.2018.2524

Rajagopalan, R., Litvan, I., & Jung, T.-P. (2017). Fall Prediction and Prevention Systems: Recent Trends, Challenges, and Future Research Directions. Sensors (Basel, Switzerland), 17(11). 10.3390/s17112509

Saftari, L. N., & Kwon, O.-S. (2018). Ageing vision and falls: a review. Journal of Physiological Anthropology, 37(1), 11. 10.1186/s40101-018-0170-1

Simôes, W. C. S. S., & De Lucena, V. F. (2016). Blind user wearable audio assistance for indoor navigation based on visual markers and ultrasonic obstacle detection. 2016 IEEE International Conference on Consumer Electronics (ICCE), 60–63. 10.1109/ICCE.2016.7430522

Studenski, S., Perera, S., Patel, K., Rosano, C., Faulkner, K., Inzitari, M., Brach, J., Chandler, J., Cawthon, P., Connor, E. B., Nevitt, M., Visser, M., Kritchevsky, S., Badinelli, S., Harris, T., Newman, A. B., Cauley, J., Ferrucci, L., & Guralnik, J. (2011). Gait speed and survival in older adults. JAMA, 305(1), 50–58. 10.1001/jama.2010.1923

Talbot, L. A., Musiol, R. J., Witham, E. K., & Metter, E. J. (2005). Falls in young, middle-aged and older community dwelling adults: perceived cause, environmental factors and injury. BMC Public Health, 5(1), 1–9. 10.1186/1471-2458-5-86

The Vision Council. (2021). The Vision Council’s organization overview sheet. https://thevisioncouncil.org/sites/default/files/assets/media/TVC_OrgOverview_sheet_2021.pdf

Treisman, A. M., & Gelade, G. (1980). A feature-integration theory of attention. Cognitive Psychology, 12(1), 97–136. http://www.ncbi.nlm.nih.gov/pubmed/7351125

Villarroel, M. A. (2021). QuickStats: Percentage of Children Aged 2–17 Years Who Wear Glasses or Contact Lenses, by Sex and Age Group — National Health Interview Survey, United States, 2019. MMWR. Morbidity and Mortality Weekly Report, 70(23), 865–865. 10.15585/MMWR.MM7023A4

Wagenmakers, E.-J., Love, J., Marsman, M., Jamil, T., Ly, A., Verhagen, J., Selker, R., Gronau, Q. F., Dropmann, D., Boutin, B., Meerhoff, F., Knight, P., Raj, A., van Kesteren, E.-J., van Doorn, J., Šmíra, M., Epskamp, S., Etz, A., Matzke, D., … Morey, R. D. (2018). Bayesian inference for psychology. Part II: Example applications with JASP. Psychonomic Bulletin & Review, 25(1), 58–76. 10.3758/s13423-017-1323-7

WHO. (2021). Falls. https://www.who.int/news-room/fact-sheets/detail/falls

Wolfe, J. M., Cave, K. R., & Franzel, S. L. (1989). Guided search: an alternative to the feature integration model for visual search. Journal of Experimental Psychology. Human Perception and Performance, 15(3), 419–433. http://www.ncbi.nlm.nih.gov/pubmed/2527952

